# Preclinical Results: Canine Phase I Safety Study of CM-101, a Tumor Capillary Specific Streptococcal Polysaccharide Toxin, for application in Spontaneous Canine Cancer

**DOI:** 10.1101/2024.03.12.584502

**Authors:** Rhett W Stout, Bonnie Boudreaux, I Horia Inegulescu, Roger A Laine

## Abstract

**Background:** The study purpose was to evaluate canine safety of CM101, a polysaccharide Group B *Streptococcus agalactiae* tumor hemorrhagic toxin therapeutic.

**Hypothesis:** CM101 specifically targets tumor vasculature as published in a human Phase 1 safety study that showed a wide therapeutic window. The hypothesis is that dogs should display a similar safety profile with low side-effects for CM101 canine cancer therapeutics.

**Animals:** Considering the previous human safety trial, and in the interest of conserving purpose-bred dogs, on advice of USDA staff, we only used two healthy males, ∼20 months old.

**Methods:** USDA advice was to administer 10x the unit dose of 7.5µg/kg to 2 dogs and if no side effects, proceed to a pilot phase II. Given the dose was 10X the effective unit dose in humans, a further dose escalation was not considered necessary. Dogs were given 10 units (75 µg/kg) CM101 in normal saline over 22 minutes intravenously. Blood and Urine were collected before infusion, intervals post infusion, and 2 weeks after. Under anesthesia through recovery, rectal temperature, heart rate and indwelling arterial blood pressure vital signs were monitored electronically. Clinical observations recorded through two weeks after infusion.

**Results:** Total WBC (white blood cell) counts dropped below normal range two hours post-infusion, after 6-11 hours rising above the normal range, returning to baseline at 52 hours post-infusion. Creatinine kinase was elevated two hours post infusion returning to baseline in 6-72 hours. Urinalysis remained within normal limits.

**Conclusions and clinical importance:** No adverse effects were observed when healthy dogs were given 10 units CM101. These finding suggest a wide therapeutic window for investigation in canine cancer.

## Background

In the 1980’s Hellerqvist, Sundell, *et al*. isolated the active polysaccharide toxin principle from Group B *Streptococcus* (GBS), a causative agent of “Early Onset Disease” in humans(Parker 1977) (Parker 1977, Hellerqvist, Rojas et al. 1981, Rojas, Green et al. 1981). This disease is seen in septic infants within five days post-partum when infected with *S. agalactiae*. The toxin largely targets lungs and peripheral capillaries leading to death in up to 30 % of untreated cases (Knox 1979). They isolated a 270kDa polysaccharide fraction from the GBS culture filtrate, characterized as protein free, LPS free and, with a lipid and phosphate attached (Hellerqvist, Rojas et al. 1981, Rojas, Green et al. 1981). Eventually they were able to identify the same toxin in urine and cerebrospinal fluid from “Early Onset Disease” sick GBS infected human neonates (Sundell, Yan et al. 2000). Because the toxin affected hypoxic driven new capillaries, it was investigated as a potential anticancer compound. Rodent studies indicated significant antitumor activity in both short and long term studies (Hellerqvist, Thurman et al. 1993, Thurman, Russel et al. 1994). This GBS toxin fraction, called CM101, was assigned IND4578 by the FDA and was evaluated in a Phase I clinical trial in volunteer stage 4 cancer patients (Quinn, Thurman et al. 1995, DeVore, Hellerqvist et al. 1997). One third of patients in the trial demonstrated tumor reduction or stabilization after one cycle CM101 treatment. Elevation of soluble e-selectin indicated capillary endothelium was the target of the induced inflammatory cascade, within the neoplastic tissue (DeVore, Hellerqvist et al. 1997, Wamil, Thurman et al. 1997).

Spontaneous cancer is common in pets. Dogs are being treated with human cancer therapeutics aggressively. Many veterinary colleges and some private practices offer sophisticated chemotherapeutic and surgical options, and various forms of radiation. To move CM101 into clinical animal trials with spontaneous cancer we needed to demonstrate safety in normal animals. Herein we report results from a Phase I safety trial of CM101 in healthy dogs.

## Materials and Methods

### Animals

The trial was approved by the University Institutional Animal Care and Use Committee. The university is fully accredited by AAALAC, International and has an approved assurance statement (#A3612-01) on file with the Office for Laboratory Animal Welfare.

Two, purpose bred, male, intact, beagle dogs approximately 20 months of age were used. Both dogs were current on vaccinations, were on heart worm preventative, and weighed approximately 13 kg each. Prior complete blood counts (CBC), chemistry panels (Chem), and physical exams (PE) indicated the dogs were in good health at the start of the study. The dogs were housed in climate controlled kennels and had *ad libitum* access to water and feed (Prolab^®^ Canine 1600, PMI^®^ Nutrition International, Inc. Brentwood MO).

### CM101 preparation and dosage determination

In the original human trial one unit of CM101 activity was delivered by a dosage of 7.5 ug/kg (DeVore, Hellerqvist et al. 1997). This unit of activity was determined in a sheep model at Vanderbilt, (Hellerqvist, Rojas, et al. 1981) which we repeated with the same results prior to this study (data not shown). In conference with the USDA it was suggested to test CM101 at 10x the starting dose used in the human clinical trial where a maximum tolerated dose (MTD) of 5 units (37.5µg/kg) was determined in stage 4 cancer patients. Thus, the test dosage of CM101 was 75 ug/kg of body weight. The total CM101 dose for dog 1 (13 kg) was 975 ug while the total dose for dog 2 (13.1 kg) was 982.5 ug.

CM101 preparation: Autoclaved culture supernatants from *Streptococcus agalactica* were made 70% in ethanol. The precipitate was subjected to phenol/water extraction, and the water phase was dialyzed against water and applied to a DE 52 column (Whatman) developed in water. The material eluted at 0.15-0.25 M NaCl was pooled, dialyzed, lyophilized, and subjected to gel filtration on a Sephacel S-300 column (5 x 100 cm) (Pharmacia). The column was developed with 0.15 M NaCl buffered at pH 6.5 with 0.01 M NaOAc. GBS toxin was collected in the void-volume peak. This partially purified GBS toxin, described previously (Hellerqvist et al. 1987), was further purified by lentil lectin chromatography. The flow-through component was dialyzed, lyophilized and subjected to HPLC gel filtration employing an Ultragel1000 column (Waters 600-MS). The column was developed in an ammonium acetate buffer and the GBS toxin eluted as an included homogeneous narrow peak, which was detected by refractive index (Waters 410). The toxin chromatographs with dextran 70 (70 kDa) and dextran 300 (298 kDa) to suggest a molecular mass of 280 kDa. The product and 150µg were aliquoted to vials and lyophilized.

Each lyophilized CM101 vial contained 150 ug of CM101. Sterile water (4.5 mls) for injection was added to a vial at room temperature. This was vortexed for 10 seconds at high speed followed by the addition of 1.5 mls of EtOH. This was vortexed for another 10 seconds yielding 25 ug/ml of CM101 in 20% EtOH.

The total dose for a given dog was drawn into a syringe with subsequent addition of 7.2% saline (0.125 mls/ml of total dose initially drawn). This solution was subsequently added to 20 mls of normal saline. Finally, additional normal saline was added to target a delivery rate of 2 mls per minute.

### Anesthesia, instrumentation, and CM101 infusion

Dogs were fasted approximately 12 hours prior to anesthesia to prevent aspiration. A given dog was sedated with Hydromorphone (Baxter Health Care Corp., Deerfield, IL) and Midazolam (Hospira, Inc., Lake Forest, IL.) intramuscularly. A cephalic catheter was placed followed by induction with propofol (Propoflo™, Abbot Laboratories, N. Chicago, IL.) intravenously to effect. Thereafter the dog was intubated and general anesthesia was maintained with isoflurane (Piramal Healthcare Limited, Andra Pradesh, India) delivered in 100% O_2_ to effect (≈2.0%). Anesthesia was maintained for 15 minutes post infusion and a given dog was recovered thereafter.

cGMP Lyophilized CM101 was provided by Carl G. Hellerqvist, Vanderbilt, Nashville, TN. The CM101 infusion was delivered *via* an infusion pump (Medex, Inc., Duluth, GA.). For a given dog, each infusion was delivered at two ml per minute for a total infusion time of 22.5 minutes each.

Antibody testing CM101: In humans, no antibodies were produced to CM101 in the Phase I trial after 12 injections over 4 weeks (Devore, et al. 1997). Sheep, mice and swine did not produce antibodies against CM101, (Hellerqvist, unpublished) possibly because the sugar polymer could not be cleaved into fragments for antigen presentation. The 2 dogs were challenged with a skin subcutaneous injection of CM101 after 2 weeks and no inflammatory response resulted concluding that, like other mammals tested, including humans, there was no immune response to the polymer.

### Data Collection

Body weight for each dog was recorded prior to anesthesia for infusion and at 2 weeks post infusion. Temperature, pulse, respiration (TPR), demeanor, and stool character were recorded at –19 hrs, 4 hrs, 6 hrs, 12 hrs, 18 hrs, 24 hrs, and at 2 weeks post infusion.

For CM101 infusion an indwelling catheter was placed in the dorsal pedal artery for blood pressure monitoring. Data were collected and monitored with two systems. Electrocardiogram (ECG), heart rate, blood pressure and temperature were recorded (1 minute prior to infusion continuously till 15 minutes after infusion) using a Powerlab 8/SP with LabChart® 7 Pro vs. 7.2 (AD Instruments, Bella Vista, NSW, Australia). ECG, heart rate, SpO_2_, respiratory rate, CO_2_ (inspired and end tidal), blood pressure and temperature were also recorded once per minute over the time frame previously stated above with a Datascope® Passport 2 with Gas Module II (Datascope Corp, Mahwah, NJ, USA).

Venous blood was collected at -19 hrs, 0 hrs, 2 hrs, 6 hrs, 11 hrs, 52 hrs, and 2 weeks post infusion. Complete blood count (CBC) parameters evaluated included erythrocyte count (RBC), hemoglobin concentration, hematocrit, red blood cell distribution width (RDW), mean corpuscular volume (MCV), mean corpuscular hemoglobin (MCH), mean corpuscular hemoglobin concentration (MCHC), platelet count, mean platelet volume (MPV), nucleated red blood cell count per 100 white blood cells (nRBC/100 WBC), plasma protein, packed cell volume (PCV), total white blood cells (TWBC’s), neutrophil count, bands, lymphocytes, monocytes, and eosinophils. The initial CBC (-19 hrs) was evaluated on a Heska CBCDiff Veterinary Hematology Sensor (Ft. Collins, CO) while all subsequent samples were analyzed on a Advia 120 (Siemens, Terrytown, NY). Chemistry panel (Chem) parameters evaluated included glucose, aspartate aminotransferase (AST), alanine aminotransferase, globulin, cholesterol, urea nitrogen, creatinine, calcium, phosphorus, sodium, potassium, chloride, bicarbonate, and anion gap. All chemistry samples were analyzed on an Olympus AU640e (Mishima Olympus Co., LTD, Japan).

Urine was collected *via* cystocentesis at 2 hrs, 11hrs, and 2 weeks post infusion and were analyzed for the following parameters: color, turbidity, specific gravity, pH, protein, glucose, ketone, bilirubin, blood hemoglobin, cast, WBCs per high power field (hpf), RBCs per hpf, and crystals.

### Data Analysis

Data are reported as descriptive statistics and results are compared to established reference ranges for dogs.

## Results

### Physical Findings

Outwardly, the dogs remained in good condition with a normal demeanor and no signs of pain or distress for the duration of the study. Similarly, no changes were noted during physical exams in body temperature, pulse, respiration, urination, stool character or quantity, or body weight for the duration of the study (data not shown). During anesthesia and infusion no abnormal changes were noted in blood pressure, ECG, heart rate, SpO_2_, inspired or end-tidal CO_2_ as well as body temperature.

### Complete Blood Count

The following CBC parameters were within normal limits for the duration of the study: RBC count, RDW, MCV, MCHC, platelets, MPV, nRBC/100 WBC, plasma protein, PCV, and eosinophils. Abnormal values were measured for hemoglobin, hematocrit, TWBC, neutrophils, bands, lymphocytes, and monocytes. Table 1 contains CBC parameters and measured values that were out of normal range, for one or both dogs, at some point during the study. Figures 1, 2, and 3 graphically illustrate the changes in TWBC, neutrophils, and lymphocytes respectively in both dogs over time.

**Table 1.** Abnormal Complete Blood Count Values Measured Before or After CM101 Administration.

| Animal ID | 1 | 2 | 1 | 2 | 1 | 2 | 1 | 2 | 1 | 2 | 1 | 2 | 1 | 2 |
| --- | --- | --- | --- | --- | --- | --- | --- | --- | --- | --- | --- | --- | --- | --- |
| Hours Relative to Infusion | -19 | -19 | 0 | 0 | 2 | 2 | 6 | 6 | 12 | 12 | 52 | 52 | 2 wks | 2 wks |
| Sample ID | 1* | 1* | 2 | 2 | 3 | 3 | 4 | 4 | 5 | 5 | 6 | 6 | 7 | 7 |
| Hemoglobin (12-18 g/dl) | 15.6 | 19.3 | 15 | 16.9 | 15.8 | 18.9 | 15.3 | 18.7 | 17 | 19 | 17.5 | 18.5 | 15.1 | 17.6 |
| Hematocrit (35-54 %) | 41.8 | 52.1 | 45.8 | 52.6 | 48.5 | 58.5 | 47.1 | 57.2 | 51 | 59.1 | 53 | 55.5 | 46 | 53.7 |
| TWBC (8-14.5 x 10<sup>3</sup>/ul) | 15.9 | 11.9 | 12.9 | 6.8 | 2.9 | 2.3 | 21.6 | 11.5 | 20.9 | 19 | 15.5 | 13.3 | 14.4 | 13.7 |
| Neutrophils (3-11.5 x 10<sup>3</sup>/ul) | 11.3 | 8.9 | 9.3 | 4.7 | 2.2 | 1.5 | 18.3 | 9.6 | 15.4 | 14 | 11.3 | 9.4 | 11.1 | 10.5 |
| Bands (0-0.3 x 10<sup>3</sup>/ul) | 0 | 0 | 0 | 0.1 | 0.3 | 0.1 | 1.7 | 1.3 | 0 | 0.2 | 0 | 0 | 0.1 | 0 |
| Lymphocytes (1-4.8 x 10<sup>3</sup>/ul) | 2.9 | 1.8 | 1.8 | 0.9 | 0.4 | 0.5 | 0.8 | 0.1 | 4.2 | 2.7 | 3.4 | 2 | 2.9 | 2.6 |
| Monocytes (0.1-1.4 x 10<sup>3</sup>/ul) | 1.4 | 1 | 1.4 | 0.8 | 0.1 | 0.1 | 0.8 | 0.5 | 1 | 1.9 | 0.6 | 1.5 | 0.3 | 0.4 |

Ten units of CM101 was delivered intravenously to healthy dogs. Total white blood cells (TWBC), weeks (wks), shaded values are outside the normal range (reference?). Samples with \* were evaluated on the Heska instrument. All others were evaluated on the Advia instrument. Normal range values are for the Advia instrument. The TWBC count for dog 1, sample 1, is within the normal range of the Heska instrument.

**Fig 1.**
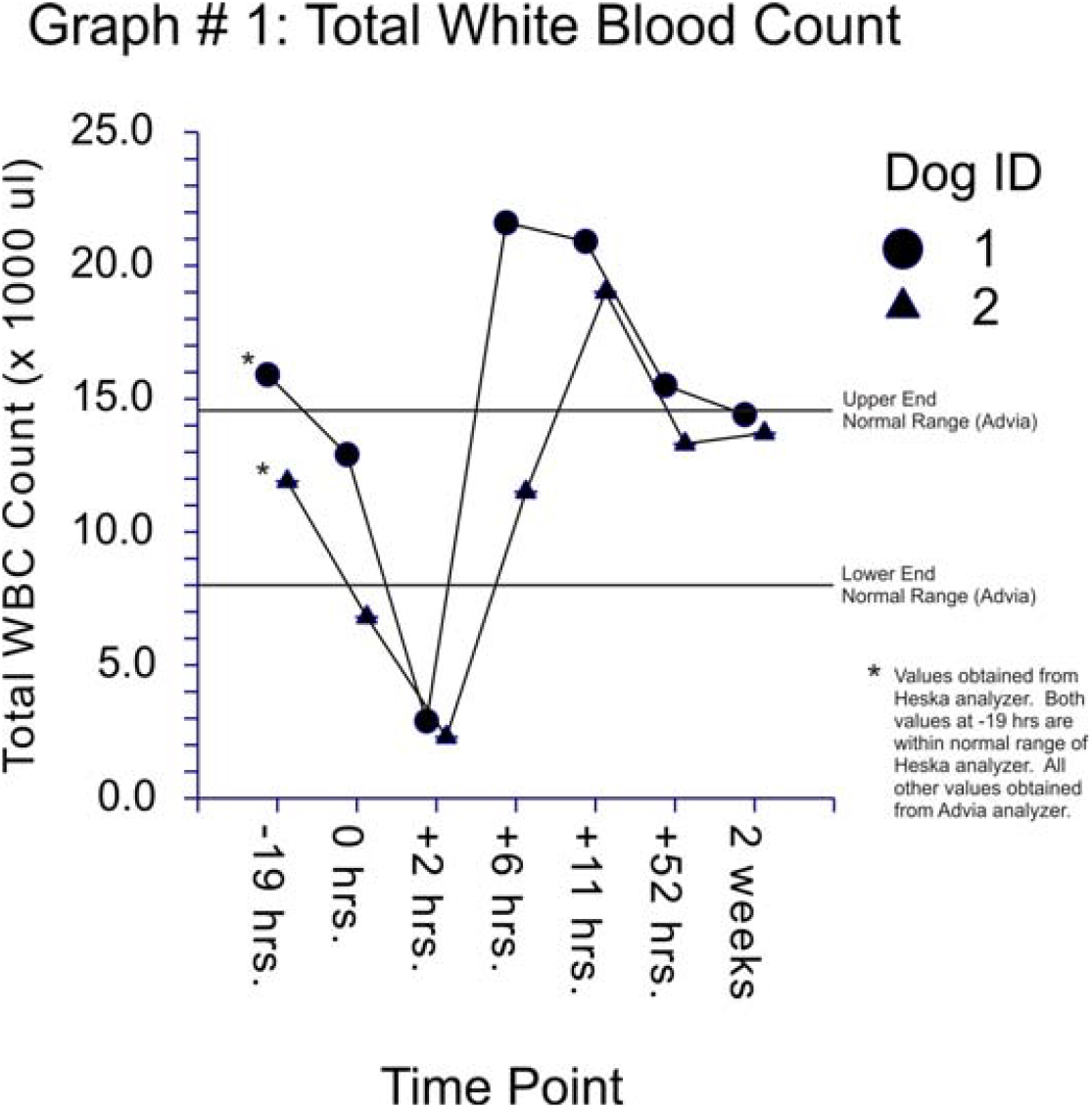
(Graph #1): Total white blood cell count before and after CM101 infusion in two dogs over time. CM101 (10 units) infusions started at 0 hours and total infusion time was approximately 22 minutes. Values with an asterisk were measured on the Heska blood analyzer. All other values were obtained using the Advia analyzer. Upper and lower normal range lines apply for the Advia analyzer. Both measurements obtained at -19 hrs were within the normal range of the Heska analyzer.

**Fig 2.**
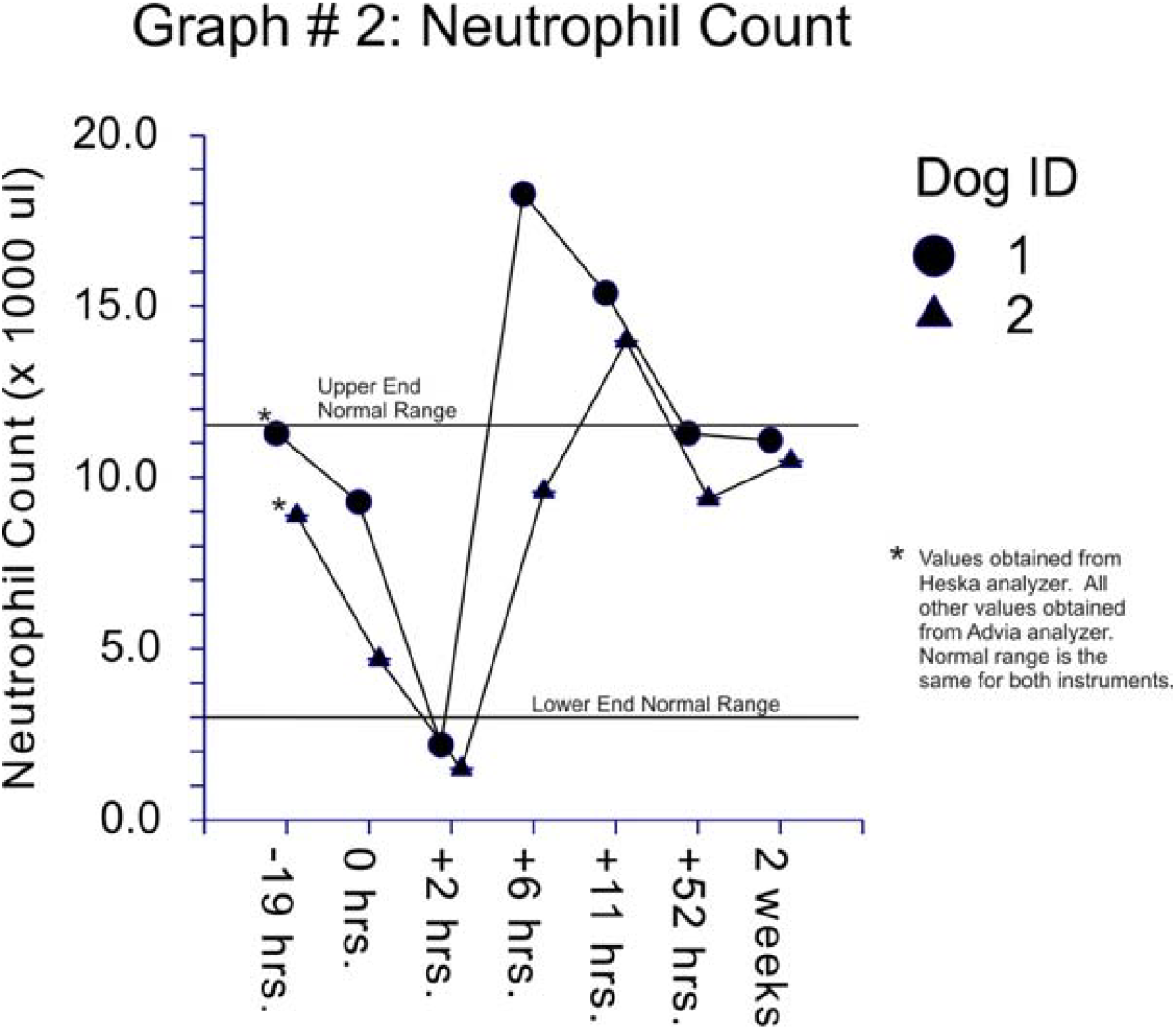
(Graph #2): Neutrophil count before and after CM101 infusion in two dogs over time. CM101 (10 units) infusions started at 0 hours and total infusion time was approximately 22 minutes. Values with an asterisk were measured on the Heska blood analyzer. All other values were obtained using the Advia analyzer. The normal range is the same for both instruments.

**Fig 3.**
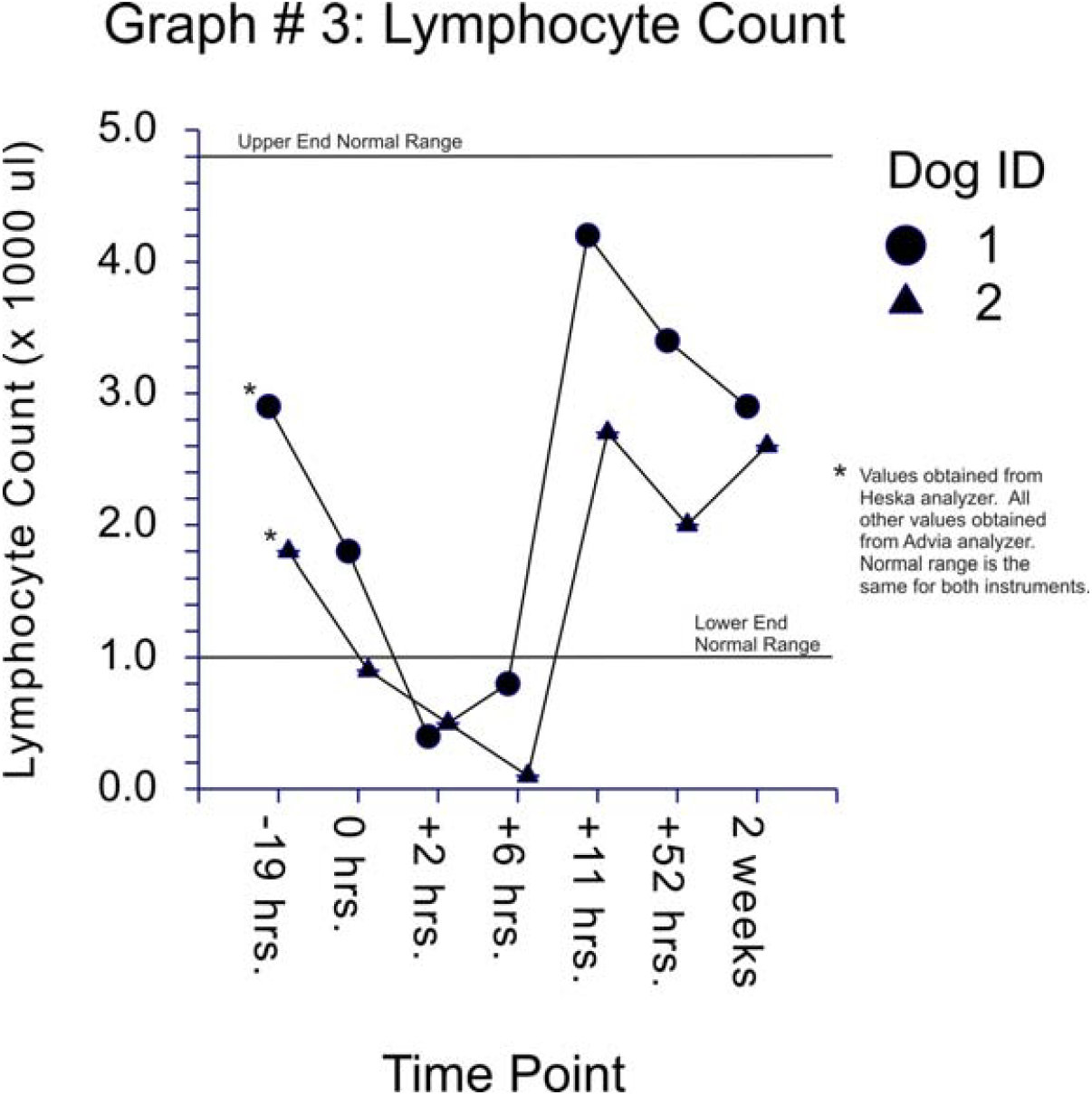
(Graph #3): Lymphocyte count before and after CM101 infusion in two dogs over time. CM101 (10 units) infusions started at 0 hours and total infusion time was approximately 22 minutes. Values with an asterisk were measured on the Heska blood analyzer. All other values were obtained using the Advia analyzer. The normal range is the same for both instruments.

### Clinical Chemistry

The following clinical chemistry parameters were within normal limits for the duration of the study: Aspartate aminotransferase, alanine aminotransferase, total protein, albumin, globulins, cholesterol, urea nitrogen, creatinine, calcium, phosphorus, sodium, potassium, chloride, bicarbonate, and anion gap (data not shown). Abnormal values were measured for glucose, alkaline phosphatase, total bilirubin, and creatinine kinase. Table 2 contains clinical chemistry parameters and measured values that were out of normal range, for one or both dogs, at some point during the study. Figures 4 and 5 graphically illustrate the changes in alkaline phosphatase and creatine kinase in both dogs over time.

**Table 2.**
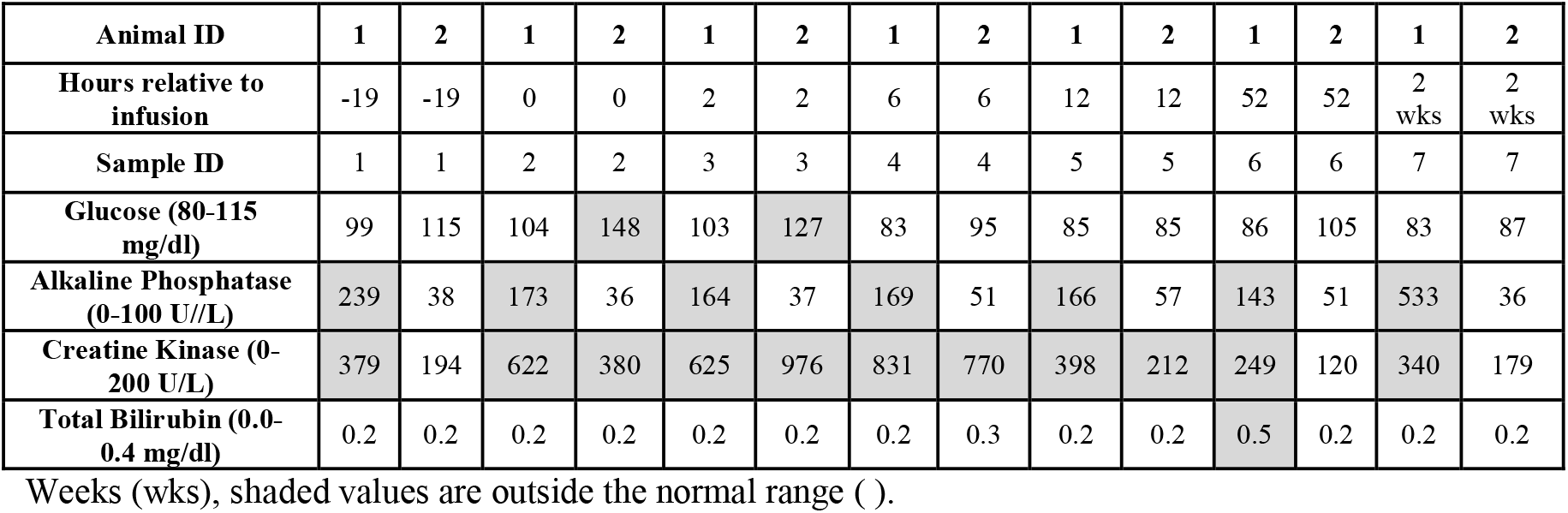
Abnormal Clinical Chemistry Values Measured Before or After CM101 Administration.

**Fig 4.**
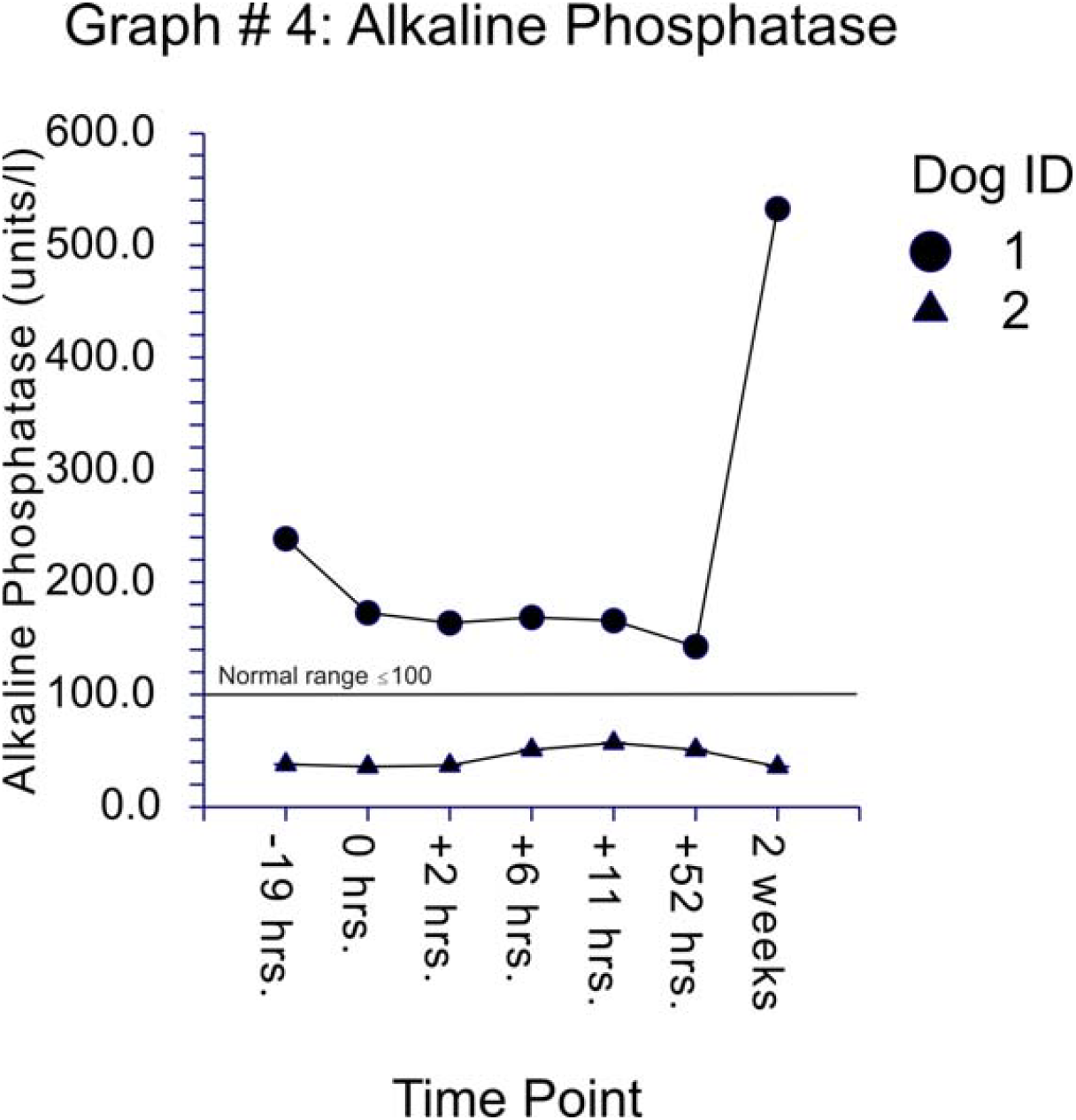
(Graph #4): Alkaline phosphatase values measured before and after CM101 infusion in two dogs over time. CM101 (10 units) infusions started at 0 hours and total infusion time was approximately 22 minutes. All samples were run on an Olympus AU640e (Mishima Olympus Co., LTD, Japan) blood chemistry analyzer. The upper limit of the normal alkaline phosphatase range for dogs was inserted into the graph for reference.

**Fig 5.**
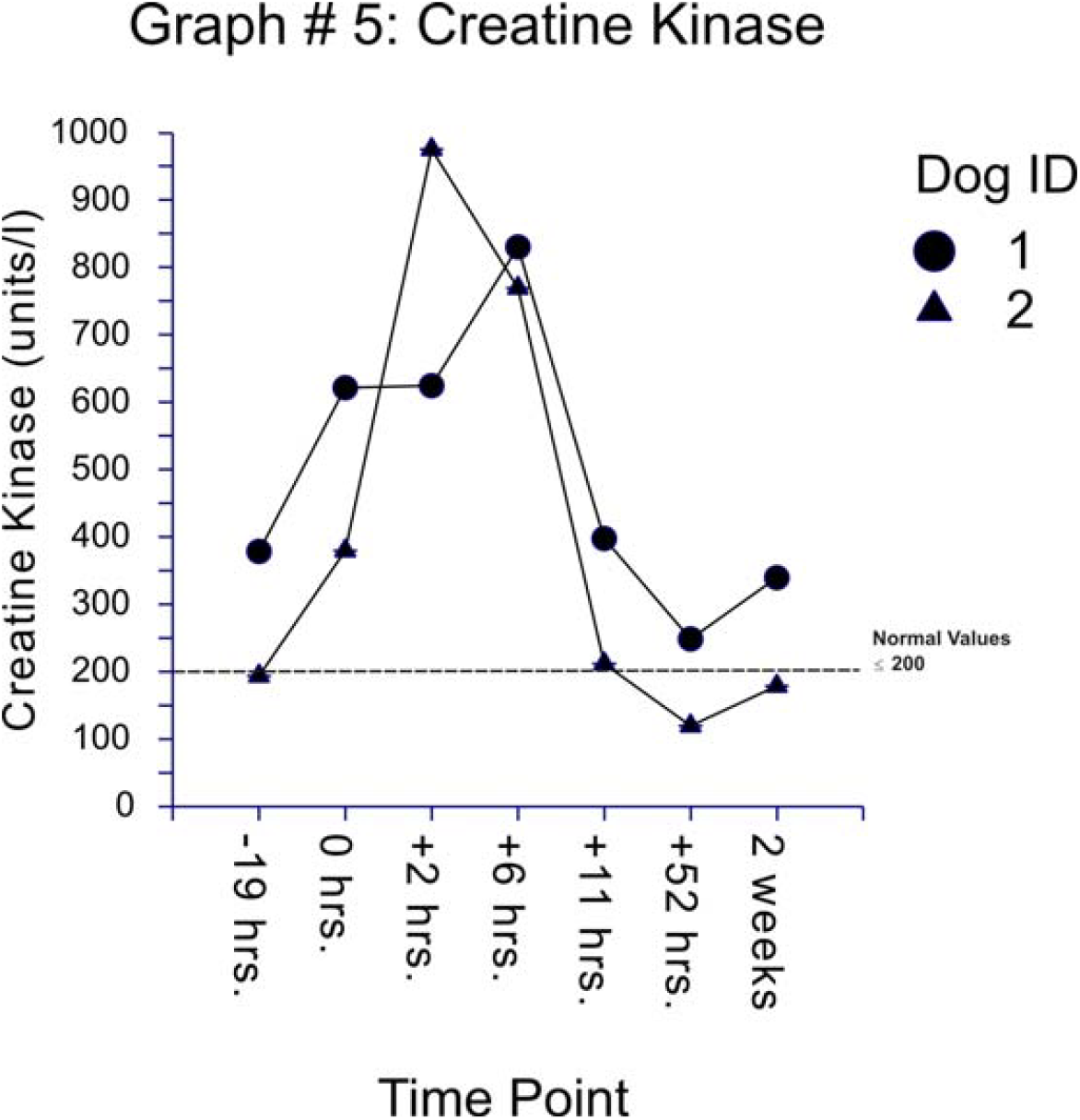
(Graph #5): Creatine kinase values measured before and after CM101 infusion in two dogs over time. CM101 (10 units) infusions started at 0 hours and total infusion time was approximately 22 minutes. All samples were run on an Olympus AU640e (Mishima Olympus Co., LTD, Japan) blood chemistry analyzer. The upper limit of the normal creatine kinase range for dogs was inserted into the graph for reference.

### Urinalysis

Urinalysis revealed some modest abnormal values over time and all measured values are provided in Table 3. Abnormal values were detected for turbidity, specific gravity, protein, bilirubin, blood hemoglobin, WBCs, and RBCs. We were unable to obtain urine from dog number 2 prior to the start of infusion.

**Table 3.**
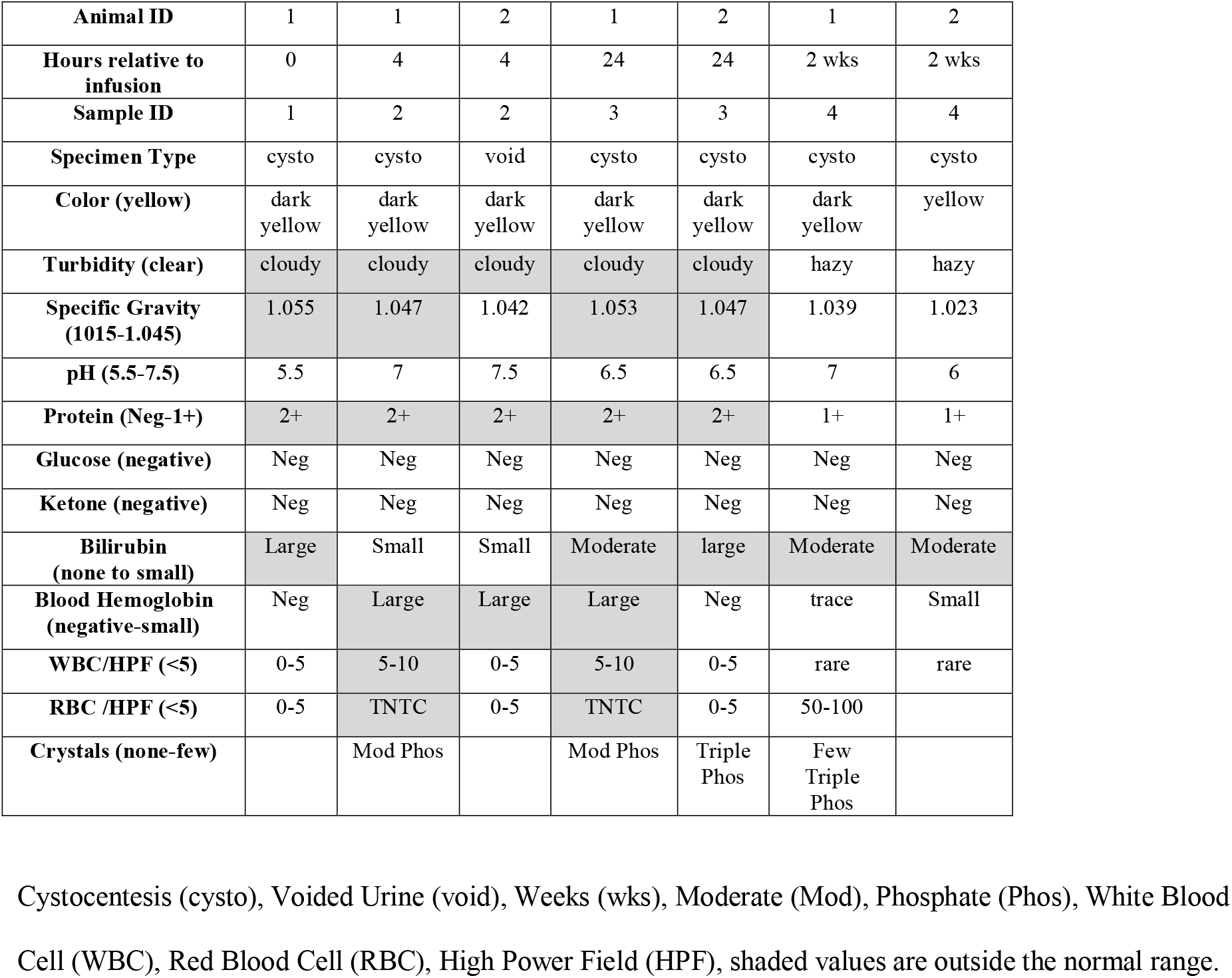
Urinalysis Values Measured at the Start of or After CM101 Administration.

## Discussion

CM101’s mode of action is not known in dogs with cancer but we presume it will be similar to that seen in other species. In human infants (≈5 days post-partum) and sheep (of any age) the toxin seemingly binds to a specific receptor present in the lung capillaries triggering its associated inflammatory response (Rojas, Green et al. 1981, Sundell, Yan et al. 2000). The inflammation appears to be a complement mediated response inducing fever, chills, depressed WBC count and with hemorrhage and necrosis in the tumor (DeVore, Hellerqvist et al. 1997, Wamil, Thurman et al. 1997). It appears that the vasculature induced by cancer cells expresses a very similar receptor (if not identical) to that which is found in the lungs of sheep and human neonates (Fu, Bardhan et al. 2001). As such, CM101’s potential to treat neoplasia has been investigated in both murine cancer models and human cancer patients with some success (Hellerqvist, Thurman et al. 1993, Thurman, Page et al. 1996, DeVore, Hellerqvist et al. 1997).

Throughout these studies authors have attempted to succinctly describe the effect of CM101 on tumor vasculature including “neovascularitis” (Thurman, Russel et al. 1994), “antineovascular” (DeVore, Hellerqvist et al. 1997) and “anti-pathoangiogenic” (Yan, Carter et al. 1998). All of these terms appear applicable, particularly within the context of each citation, but to clarify CM101 *does not prevent angiogenesis in tumors but instead interacts with new vasculature*, within a given tumor, expressing the proper receptor. There is another class of tumor therapeutic which CM101 has never been associated with but is currently receiving considerable interest in research and for clinical application. These agents are called “vascular targeting agents” (Zhang and Harris 1998) or “vascular disrupting agents” (Siemann and Horsman 2004). Given the proposed mechanism CM101 it appears this toxin derived from GBS certainly could be considered a vascular disrupting agent.

While CM01’s identification, along with its effects in neonates and tumors, were first described by Hellerqvist *et al*. we suggest this toxin produced by GBS may actually be one of the active components, if not the primary active component, of a historical cancer therapy, that being Coley’s Toxin. Coley, a surgeon, treated numerous types of cancer around the turn of the 20^th^ century with various combinations of *Streptococcus* (reported as *Streptococcus* of erysipelas) and *Serratia marcescens* (then known as *Bacillus prodigiosus*) with some notable successes (Coley 1893, Coley 1896). Through the course of numerous patients and various combinations of the previously mentioned organisms, or heat killed filtrates thereof, Coley concluded that toxins produced by these bacteria were responsible for the effects seen in the tumors, not the bacteria themselves. Erysipelas technically is a clinical diagnosis with specific clinical skin rash manifestations, yet can be caused by several different organisms. Most authors agree that *Strep. pyogenes* is the primary cause of Erysipelas yet there appears to be some debate on the next most common etiologic agent. Regardless, *Strep. agalactiae* is often included in the list of causes (Binnick, Klein et al. 1980, Blackberg, Trell et al. 2015, Stevens and Bryant 2016). If CM101 is one of the active components of Coley’s Toxin we’re left with two questions: 1. What is the true identity of the *Streptococcus* that was used to make Coley’s Toxin? 2. If Coley’s Toxin was derived, in part, from *Strep. pyogenes* and not *Strep agalactiae*, is it possible these two species, or strains thereof, share a common toxin? Literature leads one to infer that the streptococcal species used to make Coley’s toxin was *Strep. pyogenes*, yet we can find no confirmation of this fact. If in fact, *Strep. pyogenes* and *Strep. agalactiae* do share a common toxic component then the true identity (Lancefield group, species, strain, etc.) of the streptococcus used by Dr. Coley may be more important for historical purposes, than of practical interest. These questions certainly deserve further research.

Results of this trial suggest CM101 is safe for use in healthy dogs, even at high doses. The dose used in our trial was 10x the putative starting dose that might be used to treat humans or animals with cancer. In the single human clinical trial reported doses range from a single unit up to 5 units of activity, the latter of which appeared to be a reasonable estimate of the maximum tolerated dose (MTD) in humans with advanced cancer (DeVore, Hellerqvist et al. 1997). At this juncture we can make no estimate of the MTD in canines and at least in healthy dogs it appears the MTD was not reached. It is reasonable to speculate the MTD in dogs with cancer will be lower than 10 units/kg assuming a robust cytokine cascade is promulgated, similar to that seen in humans (DeVore, Hellerqvist et al. 1997, Wamil, Thurman et al. 1997). In the future we plan to measure cytokines in our experiments which likely would help to complete the picture of how CM101 would work in dogs.

Clinically speaking no toxic events were observed for the duration of the trial and had we not performed routine blood work, along with urinalysis, there would be little to discuss concerning effects in healthy dogs. Data presented in Table 1 suggest the dogs were slightly dehydrated with mild elevations in hematocrit. We suspect dehydration resulted from fasting prior to infusion with CM101. Fasting dogs is routine prior to general anesthesia to prevent aspiration of stomach contents should the animal vomit. Water is typically not withheld until the morning of the anesthetic event (as was the case for this trial), however we did not know the last time the animals drank prior to water removal early on the day of infusion. It is doubtful clinical use of CM101 would require anesthesia in most patients. Anesthesia was used in our trial to maximize data collection during the infusion without the risk of leads or catheters being removed by a rambunctious patient. Ethically, it was felt that the anesthetic would have minimal effects on the CM101 effects, and that a conscious dog with the leads, IV, monitors would have needed to be extensively restrained during the procedure, and be made extremely uncomfortable.

Figures 1-3 illustrates the most interesting results elucidated from the CBC data, perhaps from the trial as a whole. Our results suggest CM101 is recruiting WBCs, primarily neutrophils, yet no fever is induced. All three figures demonstrate similar responses with a precipitous drop in cell counts, below normal range, with a nadir at approximately two hours. This is followed by a robust rebound in relatively short order, with a subsequent fall back towards and finally into the normal range. Interestingly, the leukocyte responses in our study are similar to findings in sheep which express the receptor for CM101 in their lungs throughout life (Rojas, Larsson et al. 1983, Sandberg, Engelhardt et al. 1987). The blood sampling number and frequency were based on measurements taken in the published human trial (Devore, et al, 1997), which showed no problems over time except dyspnea in patients with lung tumor involvement, however it should be noted that the human trial comprised stage 4 cancer patients. We suspect had we sampled blood more often the timing pattern would have more closely matched results reported in sheep. We repeated the sheep test (Rojas, Larsson et al. 1983, Sandberg, Engelhardt et al. 1987) 5 times to ensure safety of the procedure before commencing the dog study data not shown, the sheep were restrained. A similar pattern was also seen with eosinophils and monocytes in our study with a precipitous drop in absolute cell numbers during the same time frame followed by a rebound although measured values largely remained within the normal range (eosinophil data not shown). Taking all leukocytes into consideration, a brief panleukopenic response was seen shortly after CM101 administration followed by a strong rebound above baseline, with neutrophils being the predominate responding cell type.

Creatine kinase is a cytosolic enzyme found primarily in skeletal and cardiac muscle in the canine with minor contributions from the brain and smooth muscle (Stockham and Scott 2008). While there are many diseases or disorders that are capable of inducing elevations of CK there are only four which might explain the elevations we recorded during this trial: anesthesia/hypotension, exercise, trauma from catheter placement or toxicity due to CM101. The maximum values measured were almost 4x the upper limit of the reference range however this level of increase would be considered mild at best (personal communication, S. Gaunt). While it has been shown that exercise in dogs does increase CK values, the amount of increase is much less than our observations with levels never reaching twice their baseline (Chanoit, Concordet et al. 2002, Queiroz, Silva et al. 2016) thus exercise as a cause for the elevated CK is ruled out.

Various anesthetic agents have been associated with elevations in CK in several species, particularly horses. Generally speaking, it appears most of the reports a more likely associated with the anesthetic’s cardiovascular effects leading to muscle hypoperfusion and not a direct toxic insult of the anesthetic agent itself. Halothane has been associated with elevations in CK in anesthetized dogs where it was compared with propofol. (Aktas, Vinclair et al. 1997) (Natalini, Krahn et al. 2017) (Natalini, Krahn et al. 2017). Since low blood pressure was not recorded in our study and we were using isoflurane as our primary anesthetic agent, during a relatively short anesthetic event, we can reasonably conclude the elevations in CK were not the result of our use of isoflurane or general anesthesia. Intramuscular injections are also associated with elevations in CK but, as above, there is little literature to support this claim (Stockham and Scott 2008). Assuming both of these sources of error are possible, the CK values recorded at -19 and 0 hrs suggest the possibility that this source of sampling error could have occurred in our study.

Alternatively, CM101 at a very high dose may be the cause of the elevations in CK. Without further testing and isoenzyme analysis, we cannot determine if this observation is associated with skeletal muscle, cardiac muscle, or other sources of tissue. In the single human trial with CM101, clinical chemistry values were recorded and no anomalies were noted with CK, although it is unknown if this enzyme was actually measured (DeVore, Hellerqvist et al. 1997). Lipopolysaccharide (LPS), unlike CM101, is well characterized within many animal models. Despite differences between LPS and CM101, the physical and inflammatory response to CM101 injection in the sheep model paints a similar picture as that seen for LPS (Hellerqvist, Rojas et al. 1981, Rojas, Green et al. 1981, Sandberg, Engelhardt et al. 1987, Yates, Loest et al. 2011). The cytokine responses to both CM101 and LPS have been investigated and IL6 specifically has direct relevance as it relates to CK levels found in our study. IL-6 and CK were shown to be elevated in both myocardial infarction, rhabdomyolysis, and exercise in humans with the both striated and cardiac muscle containing rich sources of CK (Ikeda, Ohkawa et al. 1992, Cruickshank, Oldroyd et al. 1994, Bruunsgaard, Galbo et al. 1997, Ostrowski, Schjerling et al. 2000, Del Coso, Valero et al. 2017). Furthermore, LPS challenge was shown to elevate CK in dogs, horses, rabbits, and rats while elevations in IL-6 were only noted in rats (Pattanaik and Prasad 1998, Peiro, Barnabe et al. 2010, Shao, Qu et al. 2011, Shih, Hii et al. 2016). IL-6 was elevated in the first human trial of CM101 and in infants with GBS infections (DeVore, Hellerqvist et al. 1997, Sundell, Yan et al. 2000). While we did not measure IL-6 in our study, when our findings were compared to the previously mentioned works with CM101 and LPS, a similar CK pattern is seen which suggest a possible relationship between CM101 and elevations in CK. Thus we cannot rule out CM101, or sample related trauma, as causes for the mild elevation in CK.

Abnormalities were noted for alkaline phosphatase (ALP) in one dog during the study. Alkaline phosphatase values were within normal limits for this dog prior to the start of the study (data not shown). The elevations were mild and tapered downward through the study except or the last day of the study were a sharp spike in ALP was noted. Alkaline phosphatase is elevated in many canine diseases and is naturally elevated in young dogs due to bone growth. The age of the animals used in this study eliminates bone growth as a cause for the ALP elevations leaving us to consider cholestasis. We followed this animal over time where the ALP values falling into the normal range approximately 2 months after the end of the study. Ultrasound and liver biopsies performed shortly after the end of the study revealed no significant findings. The other dog remained in the normal range for the duration of the trial. To our knowledge ALP was not measured in any species where CM101 was tested except in the human clinical trial. In this trial ALP was elevated in 3 of their patients however, these patients had documented liver metastasis (DeVore, Hellerqvist et al. 1997). While we can’t rule out some effect of CM101 on ALP elevations in the one dog, data suggest there is no specific effect.

Largely, the urinalysis was unremarkable. Abnormal findings for urine specific gravity, protein, and bilirubin can be attributed to mild dehydration while the hemoglobin and RBC can be attributed to cystocentesis as a means of urine collection. The turbidity of the urine could be attributed to dehydration and the cellular contents of the urine sample. Interestingly, CM101 is excreted in the urine (Sundell, Yan et al. 2000). We did not analyze the dog’s urine for CM101 but perhaps the high dose of CM101 may have exacerbated the cloudiness of the urine. Future studies can examine this finding more closely.

We could have improved our study in a number of ways including measurement of cytokines, possibly increasing animal numbers, and collecting samples for analysis more frequently. Furthermore, testing canines with multiple doses over time would have allowed us to evaluate any cumulative effects of repeat dosing, especially with the changes we noted in the WBC and CK levels. All of these variables can be explored further in clinical trials.

## Conclusions

In summary, CM101 appears to be safe in healthy dogs, even at very high doses. Our findings suggest clinical trials with CM101 in canines can evaluate escalating doses of CM101 with little worry about side effects to body systems unaffected by the neoplasia at hand. Further research is needed to help clarify why the responses to CM101 in healthy dogs are so similar to patterns reported by others to LPS injection, yet no fever was recorded. Dose limiting toxicity will likely be closely tied to the size of the tumor, associated metastasis, and inflammatory cascade promulgated by the interaction of CM101 with tumor vasculature. Coley’s toxin component or not, a clinical trial is underway.

